# ERGA-BGE genome of Noah’s Ark shell (*Arca noae* Linnaeus, 1758), a Mediterranean bivalve species

**DOI:** 10.1101/2025.02.10.632159

**Authors:** Katerina Vasileiadou, Tereza Manousaki, Thanos Dailianis, Grigorios Skouradakis, Emmanouela Vernadou, Danae Karakasi, Astrid Böhne, Rita Monteiro, Rosa Fernández, Nuria Escudero, Genoscope Sequencing Team, Alice Moussy, Corinne Cruaud, Karine Labadie, Lola Demirdjian, Emilie Téodori, Simone Duprat, Patrick Wincker, Pedro H. Oliveira, Jean-Marc Aury, Francesca Floriana Tricomi, Leanne Haggerty, Fergal Martin, Chiara Bortoluzzi

## Abstract

*Arca noae*, also known as the Noah’s Ark clam, is a bivalve mollusk found in the shallow coastal waters of the Mediterranean Sea and the eastern Atlantic Ocean. This species plays a crucial ecological role by filtering plankton and organic particles from the water, helping maintain water quality and supporting nutrient cycling in marine ecosystems. It is also an important food source for various marine predators, including fish and crustaceans, thereby contributing to the coastal food web. *Arca noae* is notably resilient to environmental stressors, such as temperature fluctuations, changes in salinity, and pollution, making it a valuable model species for studying how bivalves adapt and respond to stress. While it is not commonly harvested commercially, *Arca noae* is of great interest to marine researchers due to its ability to thrive in diverse coastal habitats. The reference genome of *Arca noae* will thus provide important evolutionary insights. The entirety of the genome sequence was assembled into 19 contiguous chromosomal pseudomolecules. This chromosome-level assembly encompasses 1.5 Gb, composed of 257 contigs and 119 scaffolds, with contig and scaffold N50 values of 20.5 Mb and 84.7 Mb, respectively.

## Introduction

*Arca noae*, or Noah’s Ark shell or Καλóγνωμη in Greek, is a bivalve mollusk of the family *Arcidae*. This mollusk has a characteristic thick shell, with long and straight hinges, shortened anteriorly and elongated posteriorly. The shell’s pigmentation is white with brown stripes from the centre to the margins (Morton & Peharda, 2008). The species inhabits rocky substrates up to 100m depth in the entire Mediterranean basin, the Black Sea, and the Eastern Atlantic coasts (Skouradakis et al., 2022). Although the species is economically important as an edible bivalve, its cultivation is challenging. Therefore, the species is primarily obtained through fishing. The species was once very abundant along the Adriatic coasts until World War II. However, the introduction of destructive fishing practices, such as bottom trawling, led to a significant decline in *A. noae* populations, followed by a slow recovery (Mautner et al., 2018). *Arca noae* is not included on the IUCN Red List nor the CITES list of protected species.

*Arca noae* is an epifaunal, protandric bivalve with a slow growth rate that inhabits rocky coasts with low salinity by freshwater inflows; it lives approximately 15 years and reaches sexual maturity at 2 years of age (Peharda et al., 2006). The growth rate has been proven to be strongly related to the salinity, temperature, and chlorophyll-α concentration of the adjacent seawater.

Developing a high-quality reference genome for *A. noae* is essential for advancing our understanding of its unique genetic makeup and adaptive traits. As the species can successfully recover from mass mortality events (Župan et al., 2012), sequencing its genome will provide insights into the genetic basis of its resilience and recovery mechanisms. Additionally, due to its overexploitation, the genome of *A. noae* could be an important resource for developing marine conservation strategies, as this species can serve as a potential indicator for monitoring the health of marine ecosystems.

The generation of this reference resource was coordinated by the European Reference Genome Atlas (ERGA) initiative’s Biodiversity Genomics Europe (BGE) project, supporting ERGA’s aims of promoting transnational cooperation to promote advances in the application of genomics technologies to protect and restore biodiversity (Mazzoni et al., 2023).

## Materials & Methods

ERGA’s sequencing strategy includes Oxford Nanopore Technology (ONT) and/or Pacific Biosciences (PacBio) for long-read sequencing, along with Hi-C sequencing for chromosomal architecture, Illumina Paired-End (PE) for polishing (i.e. recommended for ONT-only assemblies), and RNA sequencing for transcriptome profiling, to facilitate genome assembly and annotation.

### Sample and Sampling Information

Katerina Vasileiadou sampled one specimen of *A. noae*, which was determined based on Manousis (2021). The specimen was identified by Katerina Vasileiadou, from the Hellenic Centre for Marine Research (Greece), on 31 May 2023. Sampling was performed under permission ΥΠΕΝ/ΔΔΔ/34284/1131 from the Ministry for Environment and Energy Secretariat General for Natural Environment & Water Directorate General for Forests & Forest Environment Directorate for Forest Management. The specimen was hand-picked in the Elounda bay, Lasithi, Crete (Greece). Tissues were removed from living specimens, flash-frozen, and preserved at -80 °C until DNA extraction.

### Vouchering information

Physical reference material for the sequenced specimen was deposited in the National History Museum of Crete https://www.nhmc.uoc.gr/ under accession number NHMC.52.27.

Frozen reference tissue material of muscle is available from a proxy individual (i.e., an individual sampled at the same time and location as the one that was sequenced) at the National History Museum of Crete https://www.nhmc.uoc.gr/ under voucher ID NHMC.52.27.

An electronic voucher image of the sequenced individual is available from ERGA’s EBI BioImageArchive dataset https://www.ebi.ac.uk/biostudies/bioimages/studies/S-BIAD1012?query=ERGA under accession ID https://ftp.ebi.ac.uk/biostudies/fire/S-BIAD/012/S-BIAD1012/Files/ERGA/SAMEA114349543_1.jpg.

### Data Availability

*Arca noae* and the related genomic study were assigned to Tree of Life ID (ToLID) ‘xbArcNoae1’ and all sample, sequence, and assembly information are available under the umbrella BioProject PRJEB77125. The sample information is available at the following BioSample accessions: SAMEA114349544 and SAMEA114349548. The genome assembly is accessible from ENA under accession number GCA_964261245.1 and- the annotated genome is available through the Ensembl Beta website (https://projects.ensembl.org/erga-bge/).

Sequencing data produced as part of this project are available from ENA at the following accessions: ERX12720997, ERX12720998, ERX12722150, ERX12722151, and ERX12733464. Documentation related to the genome assembly and curation can be found in the ERGA Assembly Report (EAR) document available at https://github.com/ERGA-consortium/EARs/tree/main/Assembly_Reports/Arca_noae/xbArcNoae1. Further details and data about the project are hosted on the ERGA portal at https://www.ebi.ac.uk/biodiversity/data_portal/44597.

### Genetic Information

The estimated genome size, based on ancestral taxa, is 1.62 Gb. This is a diploid genome with a haploid number of 14 chromosomes (2n=28) and unknown sex chromosomes. All information for this species was retrieved from Genomes on a Tree (Challis et al., 2023).

### DNA/RNA processing

DNA was extracted from 160 mg of muscle tissue using a conventional CTAB extraction followed by purification using Qiagen Genomic tips (QIAGEN, MD, USA). A detailed protocol is available at protocols.io (https://www.protocols.io/view/hmw-dna-extraction-for-long-read-sequencing-using-bp2l694yzlqe/v1). DNA fragment size selection was performed using Short Read Eliminator (PacBio, CA, USA). Quantification was performed using a Qubit dsDNA HS Assay kit (Thermo Fisher Scientific) and integrity was assessed in a FemtoPulse system (Agilent). DNA was stored at 4 °C until usage. RNA was extracted from muscle (70 mg) using the RNeasy Plus Universal kit (Qiagen) following manufacturer instructions. Residual genomic DNA was removed with 6U of TURBO DNase (2 U/μL) (Thermo Fisher Scientific).

Quantification was performed using a Qubit RNA HS Assay kit and integrity was assessed in a Bioanalyzer system (Agilent). RNA was stored at -80 °C.

### Library Preparation and Sequencing

Long-read DNA libraries were prepared with the SMRTbell prep kit 3.0 following manufacturers’ instructions and sequenced on a Revio system (PacBio). Hi-C libraries were generated from muscle tissue using the Dovetail Omni-C kit (following the Insects & marine invertebrates protocol v1.2) and sequenced on a NovaSeq 6000 instrument (Illumina) with 2×150 bp read length. Poly(A) RNA-Seq libraries were constructed using the Illumina Stranded mRNA Prep, Ligation kit (Illumina) and sequenced on an Illumina NovaSeq 6000 instrument.

In total, 48x PacBio HiFi and 24x HiC data were sequenced to generate the assembly.

### Genome Assembly Methods

The genome of *Arca noae* was assembled using the Genoscope GALOP pipeline (https://workflowhub.eu/workflows/1200). Briefly, raw PacBio HiFi reads were assembled using Hifiasm v0.19.5-r593 (Cheng et al., 2021). Retained haplotigs were removed using purge_dups v1.2.5 (Guan et al., 2020) with default parameters and the proposed cutoffs. The purged assembly was scaffolded using YaHS v1.2 (Zhou et al., 2023) and assembled scaffolds were then curated through manual inspection using PretextView v0.2.5 to remove false joins and incorporate sequences not automatically scaffolded into their respective locations within the chromosomal pseudomolecules. The mitochondrial genome was assembled as a single circular contig using Oatk v1.0 (Zhou et al., 2024) and included in the released assembly.

### Genome Annotation Methods

A gene set was generated using the Ensembl Gene Annotation system (Aken et al., 2016), primarily by aligning publicly available short-read RNA-seq data from BioSamples: SAMEA114349546 and SAMN03009424 to the genome. Gaps in the annotation were filled via protein-to-genome alignments of a select set of clade-specific proteins from UniProt (Consortium, 2019), which had experimental evidence at the protein or transcript level. At each locus, data were aggregated and consolidated, prioritising models derived from RNA-seq data, resulting in a final set of gene models and associated non-redundant transcript sets. To distinguish true isoforms from fragments, the likelihood of each open reading frame (ORF) was evaluated against known metazoan proteins. Low-quality transcript models, such as those showing evidence of fragmented ORFs, were removed. In cases where RNA-seq data were fragmented or absent, homology data were prioritised, favouring longer transcripts with strong intron support from short-read data. The resulting gene models were classified into two categories: protein-coding, and long non-coding. Models that did not overlap protein-coding genes, and were constructed from transcriptomic data were considered potential lncRNAs. Potential lncRNAs were further filtered to remove single-exon loci due to their unreliability. Putative miRNAs were predicted by performing a BLAST search of miRBase (Kozomara et al., 2019) against the genome, followed by RNAfold analysis (Gruber et al., 2008). Other small non-coding loci were identified by scanning the genome with Rfam (Kalvari et al., 2018) and passing the results through Infernal (Nawrocki & Eddy, 2013). Summary analysis of the released annotation was carried out using the ERGA-BGE Genome Report ANNOT Galaxy workflow (https://workflowhub.eu/workflows/1096).

## Results

### Genome Assembly

The genome assembly has a total length of 1,496,024,100 bp in 119 scaffolds including the mitogenome (Figures 1 & 2), with a GC content of 33.6%. The assembly has a contig N50 of 20,465,118 bp and L50 of 22 and a scaffold N50 of 84,746,044 bp and L50 of 8. The assembly has a total of 138 gaps, totalling 16.0 kb in cumulative size. The single-copy gene content analysis using the Eukaryota odb10 database with BUSCO (Manni et al., 2021) resulted in 99.2% completeness (98.4% single and 0.8% duplicated). 66.2% of reads k-mers were present in the assembly and the assembly has a base accuracy Quality Value (QV) of 60.8 as calculated by Merqury (Rhie et al., 2020).

**Figure 1.**
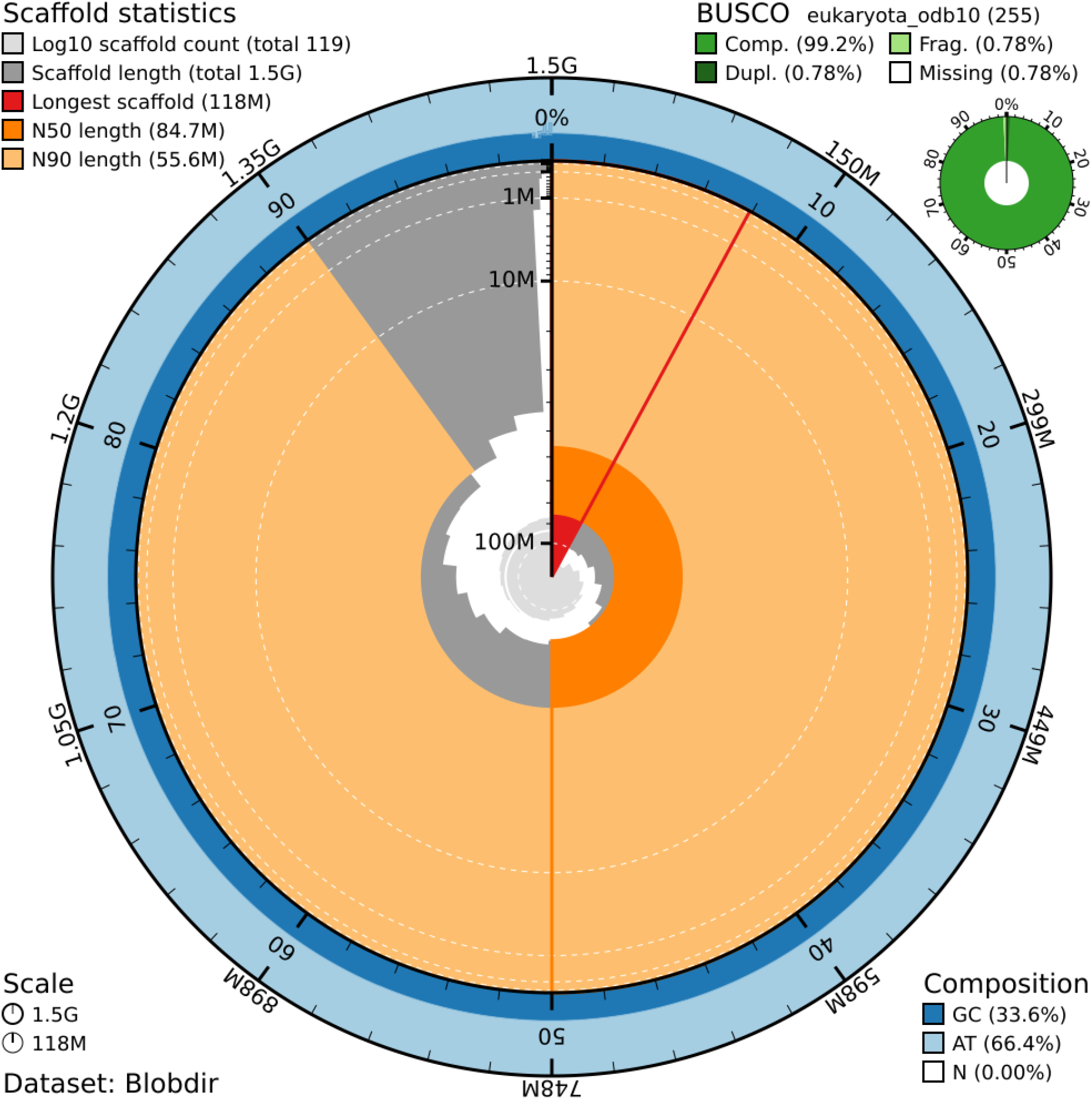
Snail plot summary of assembly statistics. The main plot is divided into 1,000 size-ordered bins around the circumference, with each bin representing 0.1% of the 1,496,024,100 bp assembly including the mitochondrial genome. The distribution of sequence lengths is shown in dark grey, with the plot radius scaled to the longest sequence present in the assembly (118 Mb, shown in red). Orange and pale-orange arcs show the scaffold N50 and N90 sequence lengths (84,746,044 bp and 55,647,231 bp), respectively. The pale grey spiral shows the cumulative sequence count on a log-scale, with white scale lines showing successive orders of magnitude. The blue and pale-blue area around the outside of the plot shows the distribution of GC, AT, and N percentages in the same bins as the inner plot. A summary of complete, fragmented, duplicated, and missing BUSCO genes found in the assembled genome from the Eukaryota database (odb10) is shown in the top right.

**Figure 2.**
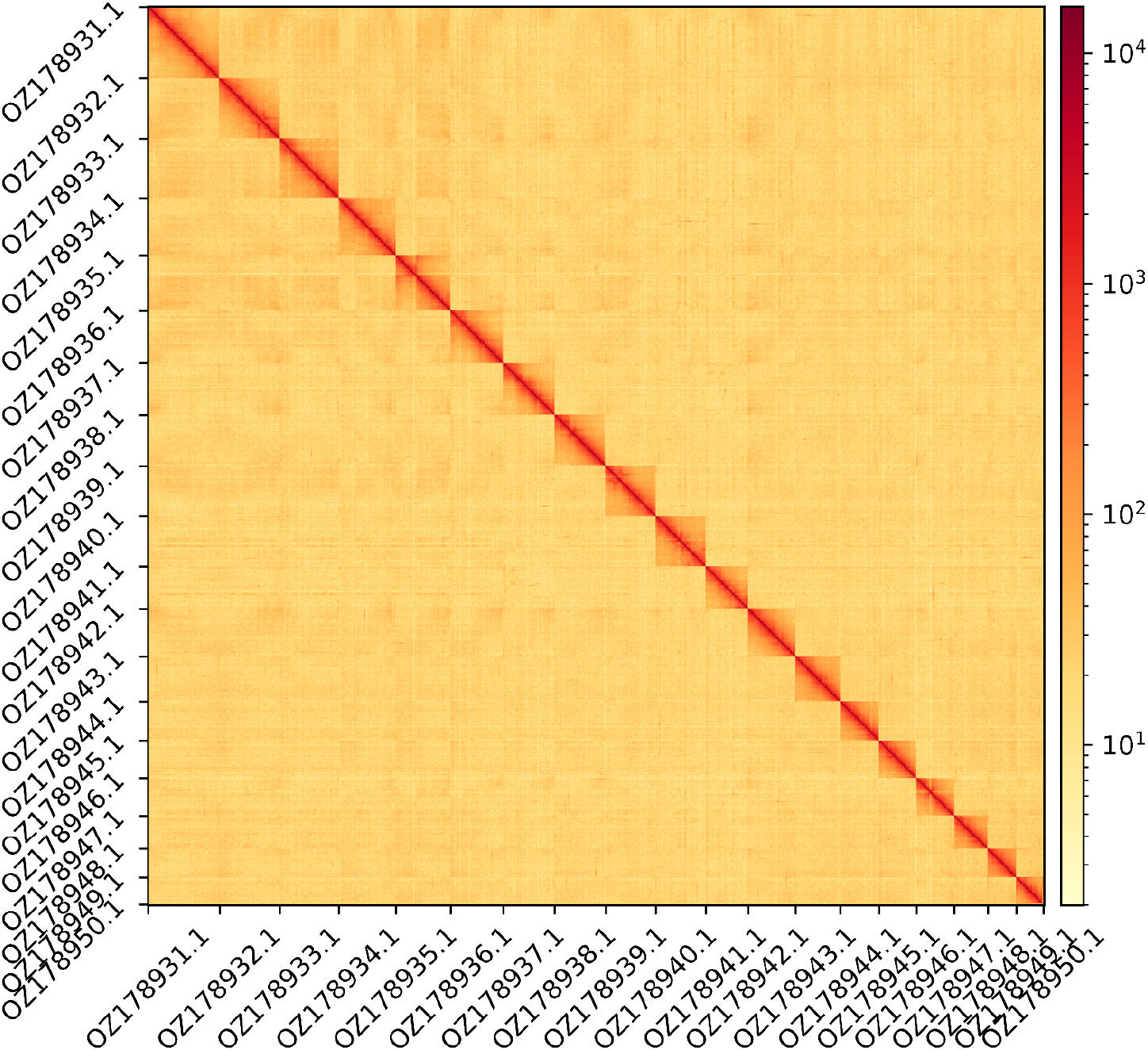
Hi-C contact map showing spatial interactions between regions of the genome. The diagonal corresponds to intra-chromosomal contacts, depicting chromosome boundaries. The frequency of contacts is shown on a logarithmic heatmap scale. Hi-C matrix bins were merged into a 25 kb bin size for plotting. Due to space constraints on the axes, only the GenBank names of the 19th largest chromosomes and the mitochondrial genome (GenBank name: OZ178950.1) are shown.

### Genome Annotation

The genome annotation consists of 17,268 protein-coding genes with associated 24,827 transcripts, in addition to 19,695 non-coding genes (Table 1). Using the longest isoform per transcript, the single-copy gene content analysis using the Eukaryota odb10 database with BUSCO resulted in 94.9% completeness. Using the OMAmer Metazoa-v2.0.0.h5 database for OMArk (Nevers et al., 2025) resulted in 9.41% completeness and 63.6% consistency (Table 2).

**Table 1.** Statistics from assembled gene models.

|  | No. genes | No. transcripts | Mean gene length (bp) | No. single-exon genes | Mean exons per transcript |
| --- | --- | --- | --- | --- | --- |
| mRNA | 17,268 | 24,827 | 21,059 | 773 | 7.2 |
| pseudogene | 0.0 | 0.0 | 0.0 | 0.0 | 0.0 |
| snoRNA | 54 | 54 | 120 | 54 | 1.0 |
| lncRNA | 4,263 | 4,630 | 8,095 | 35 | 2.5 |
| miRNA | 0.0 | 0.0 | 0.0 | 0.0 | 0.0 |
| snRNA | 366 | 366 | 145 | 366 | 1.0 |
| rRNA | 270 | 270 | 362 | 270 | 1.0 |
| scRNA | 1 | 1 | 128 | 1 | 1.0 |
| Other ncRNA | 14,741 | 14,741 | 58-70 | 14,741 | 1.0 |

**Table 2.** Annotation completeness and consistency scores calculated by BUSCO run in protein mode (eukaryota_odb10) and OMArk (Metazoa-v2.0.0.h5)

|  | Complete | Singular | Duplicated | Fragmented | Missing |
| --- | --- | --- | --- | --- | --- |
| BUSCO | 242 (94.9%) | 240 (94.1%) | 2 (0.8%) | 12 (4.7%) | 1 (0.4%) |
| OMArk | 2,026 (94.1%) | 1,865 (86.6%) | 161 (7.5%) | - | 127 (5.9%) |
|  | Consistent | Inconsistent | Contaminants | Unknown |  |
| OMArk | 10,980 (63.6%) | 1,626 (9.4%) | 0.0 (0.0%) | 4,661 (26.9%) |  |

## Acknowledgements

We would like to acknowledge the assembly reviewer, Thomas Brown, from the Leibniz Institute for Zoo and Wildlife Research (Germany). The authors acknowledge the support of the Freiburg Galaxy Team: Saim Momin and Björn Grüning, Bioinformatics, University of Freiburg (Germany), funded by the German Federal Ministry of Education and Research BMBF grant 031 A538A de.NBI-RBC and the Ministry of Science, Research and the Arts Baden-Württemberg (MWK) within the framework of LIBIS/de.NBI Freiburg.

## Conflict of Interest

The authors declare no conflict of interest related to this study. The funding sources had no involvement in the study design, collection, analysis, or interpretation of data; in the writing of the manuscript; or in the decision to submit the article for publication. All authors have participated sufficiently in the work to take public responsibility for the content and agree to the submission of this manuscript.

## Funder Information

This project received funding from Biodiversity Genomics Europe (Grant no.101059492), which is funded by Horizon Europe under the Biodiversity, Circular Economy and Environment call (REA.B.3); co-funded by the Swiss State Secretariat for Education, Research and Innovation (SERI) under contract numbers 22.00173 and 24.00054; and by the UK Research and Innovation (UKRI) under the Department for Business, Energy and Industrial Strategy’s Horizon Europe Guarantee Scheme. This work was supported by the Genoscope, the Commissariat à l’Energie Atomique et aux Énergies Alternatives (CEA), France Génomique (ANR-10-INBS-09-08) and the exploratory research programme “ATLASea: Atlas of marine genomes” and its targeted project SEQ-Sea (ANR-22-EXAT-0003-SEQ-Sea).

## Author Contributions

KV and TM coordinated the project; TD, GS and EV collected the species; TD, GS and EV identified the species; KV, TM and DK sampled and preserved biological material and provided metadata; RM, NE, RF, and AsB provided support in sampling, shipping of biological material, metadata collection, and management; the GST extracted DNA, prepared libraries, and performed sequencing under the supervision of AM, CC, KL, PHO and PW; LD, ET, SD and JMA performed genome assembly and curation; LH, SS, and FM performed genome annotation; CB generated the analysis and report. All authors contributed to the writing, review, and editing of this genome note and read and approved the final version. This work is part of the species assigned to Genoscope, which was instrumental in the wet lab, sequencing, and assembly processes, and represents a key contribution to BGE’s outputs.

## Author Information

Members of the Genoscope Sequencing Team are listed here: https://zenodo.org/records/1461149.

## Notes

https://www.ebi.ac.uk/ena/browser/view/PRJEB77125

### Competing Interest Statement

The authors have declared no competing interest.

### Summary of Updates

This version of the manuscript has been revised to update the Genome Annotation Methods.

